# Persistence and expansion of cryptic endangered red wolf genomic ancestry along the American Gulf coast

**DOI:** 10.1101/2021.04.09.439176

**Authors:** Bridgett M. vonHoldt, Kristin E. Brzeski, Matthew L. Aardema, Christopher Schell, Linda Y. Rutledge, Steven R. Fain, Amy Shutt, Anna Linderholm, William J. Murphy

## Abstract

Admixture and introgression play a critical role in adaptation and genetic rescue that has only recently gained a deeper appreciation. Here, we explored the geographic and genomic landscape of cryptic ancestry of the endangered red wolf that persists within the genome of a ubiquitous sister taxon, the coyote, all the while the red wolf has been extinct in the wild since the early 1980s. We assessed admixture across 102,621 SNP loci genotyped in 293 canid genomes. We found support for increased red wolf ancestry along an east-to-west gradient across the southern United States that was associated with historical admixture in the past 100 years. Southwestern Louisiana and southeastern Texas, the geographic zone where the last red wolves were known prior to their extinction in the wild, contained the highest and oldest levels of red wolf ancestry. X-linked regions of low recombination rates were depleted of introgression, relative to the autosomes, suggestive of the large X effect and enrichment with loci involved in maintaining reproductive isolation. Recombination rate was positively correlated with red wolf ancestry across coyote genomes, consistent with theoretical predictions. The geographic and genomic extent of cryptic red wolf ancestry can provide novel and variable genomic resources for the survival of the endangered red wolf.

## Introduction

As the field of evolutionary biology moves towards the web-of-life framework, there is broader recognition regarding the role that introgression and admixture have on adaptation and genetic rescue (Hufbauer et al. 2015; Hamilton and Miller 2016; Kronenberger et al. 2018; vonHoldt et al. 2018; Burgarella et al. 2019). However, the strength and permeability of reproductive barriers shapes the variation in both ancestry proportions and linked phenotypes that result from introgression (Haasl and Payseur 2016; Harrison and Larson 2016). Barrier loci substantially reduces gene flow between species or types, and influence the landscape of introgression across the genome (Barton and Bengtsson 1986). In species that are spatially structured, ancestry inferences are thus challenged by geographic heterogeneity, although such clines can be leveraged to explore variability in introgression rates along an admixture gradient (Gombert and Buerkle 2011). Further, recombination rates vary across the genome and can consequently result in the inference of different evolutionary histories (Lotterhos 2019). For instance, regions of low recombination have more extensive linkage disequilibrium (LD), higher differentiation, reduced nucleotide diversity, and possibly more recent coalescence relative to regions with more frequent recombination events (Geraldes et al. 2011; Seixas et al. 2018; Stevison and McGaugh 2020). However, these regions also experience more effective selection against introgression, suggested due to their linkage to and frequent enrichment with barrier loci that impact hybrid fitness (Schumer et al. 2018; Martin et al. 2019). Theory then predicts that introgression and ancestry tract lengths have a negative relationship with the time since the initial hybridization event (Pool and Nielsen 2009).

The red wolf (*Canis rufus*) has been functionally absent from the landscape since the early 1980s, yet coyotes (*C. latrans*) in southeastern Texas and southwestern Louisiana continue to harbor red wolf genetic ancestry (Heppenheimer et al. 2018a, 2020; Murphy et al. 2018). This geographic region is where the last wild red wolves were captured in the 1970s to initiate a captive breeding program as part of their Species Survival Plan (SSP) prior to their extinction in the wild (Carley 1975). Hybridization between these canid lineages prior to their captive founding has been reported (McCarley 1962; Riley & McBride 1971; Paradiso & Nowak 1972), and the persistence of canids resembling red wolf hybrids has continued to be documented through the 1990s (Giordano and Pace 2000). While initial genetic assessments of canids in Texas did not detect substantial admixture (Hailer and Leonard 2008), more comprehensive genomic surveys suggest that cryptic red wolf ancestry may be widespread in coyotes along the Gulf coast (Heppenheimer et al. 2018a, 2020; Murphy et al. 2019). Red wolves remain critically endangered with less than 10 known wild wolves persisting in a reintroduced population in North Carolina and approximately 250 in SSP captive facilities. The species suffers from reduced effective size as a consequence of founding the extant population from 14 individuals and the propensity of wild individuals to hybridize presents a significant conservation concern (Hinton et al. 2013; Brzeski et al. 2014). There is an urgent need to better understand the timing and extent of historic introgression to effectively mitigate contemporary hybridization and identify ghost red wolf genetic variants in extant coyotes which could be leverage for the future of red wolf recovery.

Genetic ancestry and timing estimates of gene flow in North American canids have been previously examined (vonHoldt et al. 2016a,b; Sinding et al. 2018). However, there is a paucity of inferences based on chromosome types (i.e. autosomal vs. sex chromosome) and integration with recombination rates. Given the predicted contrasting patterns of demographic estimates for regions of differing recombination and evolutionary rates, we hypothesized that such genomic regions may show contrasting and divergent patterns in ancestry and gene flow. We explore the impact of variable recombination rates and admixture across a dense sampling of canids from the southeastern range of coyotes and red wolves. Here, we genotyped 102,621 genome-wide SNPs in 310 canid genomes that traverse the remnant hybrid zone in southeastern Texas and southwestern Louisiana (Heppenheimer et al. 2018a, 2020; Murphy et al. 2018). Our objective was to determine if the recently discovered cryptic red wolf ancestry has a much broader geographic distribution. We were motivated to describe the extent to which undetected introgressed alleles from an endangered and extirpated species occupy the landscape. We hypothesized that introgressed red wolf ancestry found in coyotes would be highest in southeastern Texas and southwestern Louisiana, the geographic areas proximal to the source of the SSP red wolf founders (McCarley 1962; Paradiso 1968; Carley 1975; MacCarley and Carley 1979; Nowak 2002). Further, we compared patterns of ancestry on the X chromosome and autosomes, which has rarely been performed in prior studies of canid phylogeography. We further aimed to explore the relationship between admixture timing and recombination rate variation to identify gene flow events that are historic, contemporary, or are signals of incomplete lineage sorting (ILS). We finally hypothesized that younger introgressed fragments have radiated away from the hybrid zone as coyotes with admixed and introgressed genomes disperse.

## Results

### Coyotes show a west-to-east geographic gradient of probable red wolf assignments

We obtained 120,621 SNPs from restriction-site associated DNA sequence (RADseq) data genotyped in 293 canids sampled across the American south (Fig. 1A,1B). Coyotes were classified as either being sampled in their pre-1900 historic range (Arizona, California, Colorado, Kansas, Missouri, Nebraska, New Mexico, Nevada, Oklahoma, Texas, and Wyoming) or their post-1900 southeastern expansion (Alabama, Georgia, Kentucky, Louisiana, and Tennessee) (Table S1). We included representatives from North American gray wolves (*C. lupus*, including Mexican wolves *C. lupus baileyi*), eastern wolves (*C. lycaon*) from Algonquin Provincial Park, Species Survival Plan (SSP) captive red wolves, and domestic dogs (Table S1).

**Figure 1.**
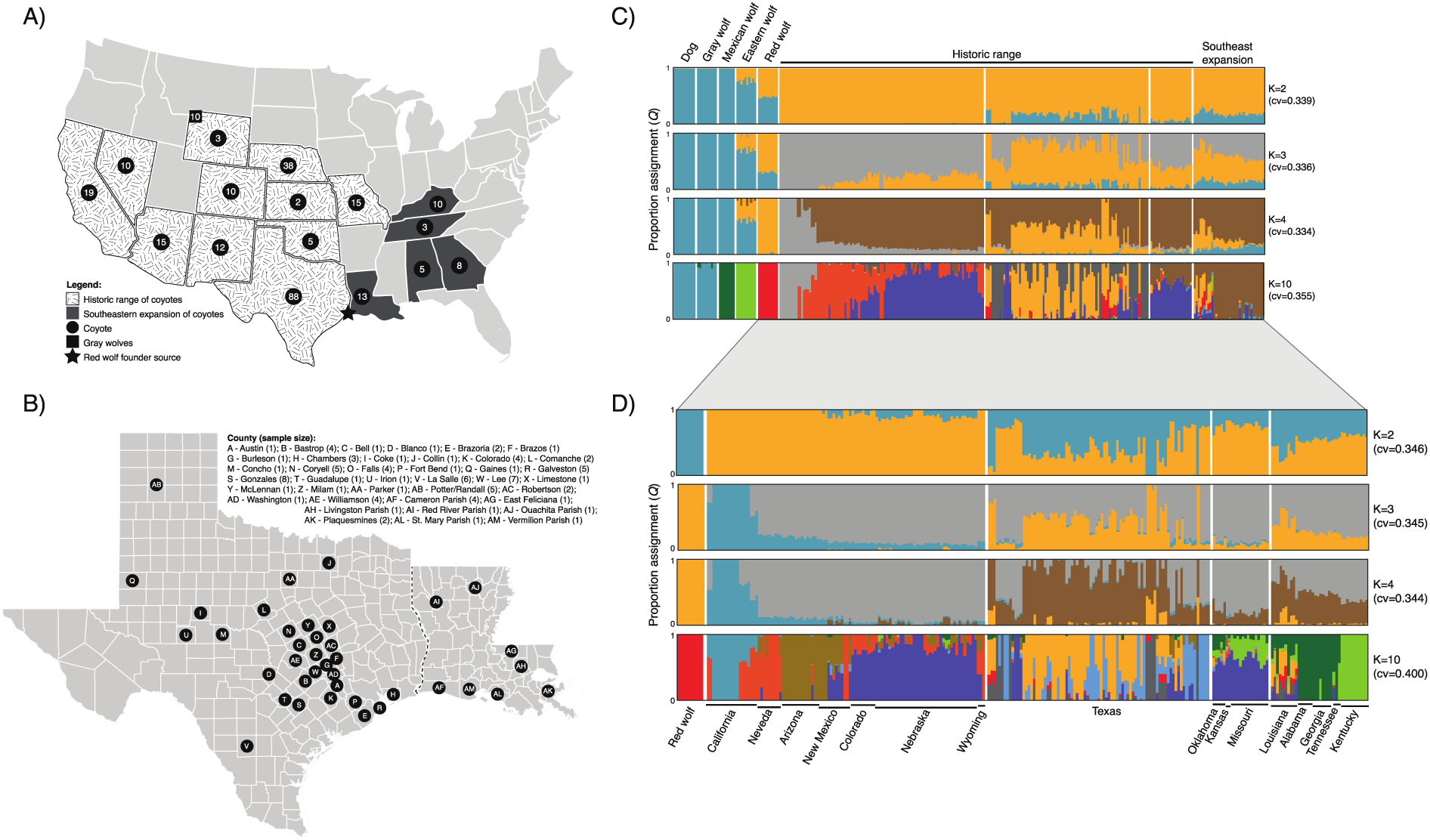
Sample map of **A)** 310 canids from the lower contiguous United States and **B)** from counties in Texas and Louisiana. Shaded states indicate either the historic or expansion range of coyotes included in this study, and numbers within symbol represent sample size. Reference genomes not included on the geographic map are: domestic dogs (n=13), eastern wolf (Algonquin Provincial Park, n=10), red wolves from the Species Survival Plan captive breeding population (n=10), and Mexican wolves from the captive breeding studbook (n=10) (see Table S1 for more details). Admixture proportions from 83K unlinked/neutral SNPs for **C)** 293 canids and **D)** 254 canids. Solid bars above or below vertical bars indicate the state from which the samples originated.

**Table 1.** *D*-statistic testing for variable P1 and P2 groups of coyotes against P3 (reference red wolves), with the outgroup of red fox. Standard errors (s.e.) and significance values were evaluated using jackknife resampling method with a block size range from 2-10Mb. No results were obtained for 2-4Mb block sizes.

|  | P1= | Historic range | Reference | Reference |
| --- | --- | --- | --- | --- |
|  | P2= | Southeastern expansion | Texas | Louisiana |
| D statistic |  | 0.156 | 0.352 | 0.369 |
| 5Mb block ($N_{\text{blocks}}=484$) | | | | |
| s.e. ($p$-value) | | 0.01 ($2.43 \times 10^{-30}$) | 0.01 ($1.85 \times 10^{-129}$) | 0.02 ($1.31 \times 10^{-49}$) |
| 6Mb block ($N_{\text{blocks}}=405$) | | | | |
| s.e. ($p$-value) | | 0.01 ($2.06 \times 10^{-28}$) | 0.02 ($8.48 \times 10^{-118}$) | 0.03 ($3.01 \times 10^{-47}$) |
| 7Mb block ($N_{\text{blocks}}=350$) | | | | |
| s.e. ($p$-value) | | 0.01 ($2.68 \times 10^{-28}$) | 0.01 ($4.91 \times 10^{-127}$) | 0.03 ($1.35 \times 10^{-47}$) |
| 8Mb block ($N_{\text{blocks}}=309$) | | | | |
| s.e. ($p$-value) | | 0.01 ($7.85 \times 10^{-26}$) | 0.02 ($8.72 \times 10^{-120}$) | 0.03 ($1.21 \times 10^{-43}$) |
| 9Mb block ($N_{\text{blocks}}=277$) | | | | |
| s.e. ($p$-value) | | 0.01 ($3.41 \times 10^{-28}$) | 0.02 ($1.21 \times 10^{-114}$) | 0.03 ($4.44 \times 10^{-44}$) |
| 10Mb block ($N_{\text{blocks}}=254$) | | | | |
| s.e. ($p$-value) | | 0.01 ($2.57 \times 10^{-27}$) | 0.01 ($4.91 \times 10^{-126}$) | 0.03 ($1.58 \times 10^{-47}$) |

A principal component analysis (PCA) of 83,851 unlinked neutral SNPs revealed a gradient in the first two PCs for coyotes, polarized by red wolves and California coyotes that explained 3.5% of the variation on PC1 (Fig. S1). This pattern was present albeit weaker in spatial distinction for the X chromosome PCA. Maximum likelihood admixture analyses across nine partitions (*K*=2-10) also supported a west-to-east geographic gradient of red wolf assignments across coyotes of the southern United States (Fig. 1C). With the inclusion of five reference groups (dogs, gray wolf, Mexican wolf, eastern wolf, and red wolf), coyotes displayed increased red wolf assignments in Texas (*K*=4 average *Q* Texas=0.42) and across the southeast expansion (Alabama=0.16, Georgia=0.14, Louisiana=0.37, Tennessee=0.10, Kentucky=0.02) relative to coyotes of the west (California=0.00, Nevada=0.00, Arizona=0.00, New Mexico=0.01) (Figs. 1C, S2; Table S2A). Red wolf assignments were refined to regions within at higher partitions (Table S2A). Although the best fit partition was *K*=2 for 830 neutral and unlinked X-linked SNPs, red wolves appeared as a distinct genetic cluster at *K*=4 (Fig. S3); as such, we found a similar trend of increased red wolf assignments in coyotes with eastward geography (Fig. S2, S3; Table S2B).

We subsequently excluded domestic dogs, eastern wolves, gray wolves, and Mexican wolves for a finer-scale assessment of 244 coyotes and 10 red wolves (254 canids sample set) across the southern United States. The PCA supported the same polarization of red wolves and western California coyotes on PC1 (2.2% variation) (Fig. S1). Admixture analysis also provided the same west-to-east gradient of increasing red wolf assignments (best fit *K*=4 average *Q* Texas=0.06, Alabama=0.03, Georgia=0.02, Louisiana=0.13, Tennessee=0.04, Kentucky=0.03) compared to western populations California=0.00, Nevada=0.00, Arizona=0.00, New Mexico=0.00) (Figs. 1D, S2; Table S2C). Higher red wolf assignments were found in Louisiana and 11 counties in southeastern Texas (Fig. S4). We focused on seven Texas counties that had a minimum of 5% probable assignment to the red wolf cluster (*K*=4 *Q* Austin=0.05, Brazoria=0.21, Brazos=0.05, Chambers=0.26, Colorado=0.12, Fort Bend=0.21, Galveston=0.43). Analysis of X-linked SNPs supported a similar pattern, albeit with greater variation in assignment values (Figs. S2, S3; Table S2D).

### Differentiating incomplete lineage sorting and gene flow

We explored the degree that incomplete lineage sorting (ILS) relative to gene flow could explain the above genetic structure patterns using the *D* statistic (Durand et al. 2011). We expanded the dataset to include data from an outgroup population comprised of data from 45 red foxes (*Vulpes vulpes*) (DeCandia et al. 2019), and repeated the SNP discovery pipeline. We catalogued 5,354,281 SNPs across 293 canids and 45 red foxes to identify derived alleles that were fixed in the red foxes as the ancestral homozygous genotype. After filtering for missing data, we retained 922,115 loci and 333 individuals; two Nebraska coyotes and three domestic dogs were excluded due to an excess of missing data. We identified a further subset of 168,004 autosomal and 1,494 X-linked loci where all 45 red foxes are fixed for a single ancestral allele, with corresponding data for the various coyote group compositions. The captive SSP red wolves (P3) were included to explore the putative source of red wolf ancestry in coyotes (P2) with respect to a reference set of coyotes as P1. We permuted the P1 and P2 groups as follows: P1=coyotes from within their historic range excluding Texas, or a reference set of coyotes identified at *K*=4 from *ADMIXTURE* as an individual that had an autosomal red wolf proportion *Q*<0.02; P2=coyotes from Texas or Louisiana (Table S2A). We found overwhelming evidence that allele sharing between coyotes and red wolves was due to gene flow and introgression rather than ILS (*D*>0.16, *p*<10x^-26^) (Table 1, Fig. S5). We found significant evidence of red wolf introgression outside the historic range of coyote, as well as in coyotes from Louisiana and Texas, corroborating past findings (e.g. Heppenheimer et al. 2018a, 2020; Murphy et al. 2018).

### Red wolf ancestry proportions are highest in southern coyote genomes

We inferred ancestry at 120K SNP loci for 138 coyotes with respect to three reference populations: gray wolves (including Mexican wolves), captive SSP red wolves, and 106 reference coyotes with negligible (*Q*<0.02) autosomal red wolf proportion at *K*=4 (Tables S2A, S3). Average red wolf ancestry proportions were highest on coyote autosomes that were collected from Texas (average=62.4%, range=8-85%) and Louisiana (average=61.4%, range=47-81%). Louisiana coyotes carried the highest average proportions of X-linked red wolf ancestry at 41% with variation across parishes (range=26-67%) (Table S3; Figs. 2, S7). At the chromosomal level, there were few regions of the genome where red wolf ancestry levels were markedly different between the Texas and Louisiana coyotes (Fig. S8). Substantial red wolf ancestry was identified in coyotes from Alabama (autosomal, X: 41%, 24%), Georgia (39%, 12%), Kentucky (23%, 4%), Missouri (15%, 4%), Oklahoma (23%, 5%), and Tennessee (34%, 28%), concordant with records of the red wolf’s historic range in these same states. We found a significant enrichment of red wolf ancestry on the autosomes relative to the X chromosome (1-tailed t-test of unequal variance, *p*=1.639−10^-20^). We documented the longest tracks of homozygous red wolf ancestry in coyotes from Alabama (autosomal, X: 3.8Mb, 3.9Mb), with longer red wolf ancestry tracks found among autosomes in Georgia (986Kb), Louisiana (403Kb), and Texas (359Kb) (Table S3). We also detected gray wolf ancestry across these coyotes (average autosomal=5.8%, X=<1%); however, due to small sample sizes, we could not determine if this was driven by Mexican wolf introgression or allele sharing with the gray wolf ancestor (Table S3).

An analysis of coyotes with county-level information for coyotes collected in Texas (74/83 individuals) and Louisiana (9/10 individuals) revealed the highest autosomal red wolf ancestry was found within counties or parishes where trapping efforts to establish the SSP red wolf founders originated, which were Jefferson and Chambers counties in Texas, and Cameron and Calcasieu parishes in Louisiana (Texas: Chambers=84%, Brazoria=83; Galveston=82%, Fort Bend=82%; Louisiana: Cameron=80%, Vermilion=78%) (Table S3; Fig. S9). We found similarly high levels of X-linked red wolf ancestry in Texas counties where some of the SSP red wolf founders were trapped: Chambers (59%), Austin (57%), and Galveston (55%), and Louisiana (Plaquemines=63%, Vermilion=52%). A noticeable gradient was observed with increasing red wolf ancestry from northwest to southeast Texas (Fig. S9), with the trend much more pronounced on the autosomes relative to the X chromosome (Fig. 2).

**Figure 2.**
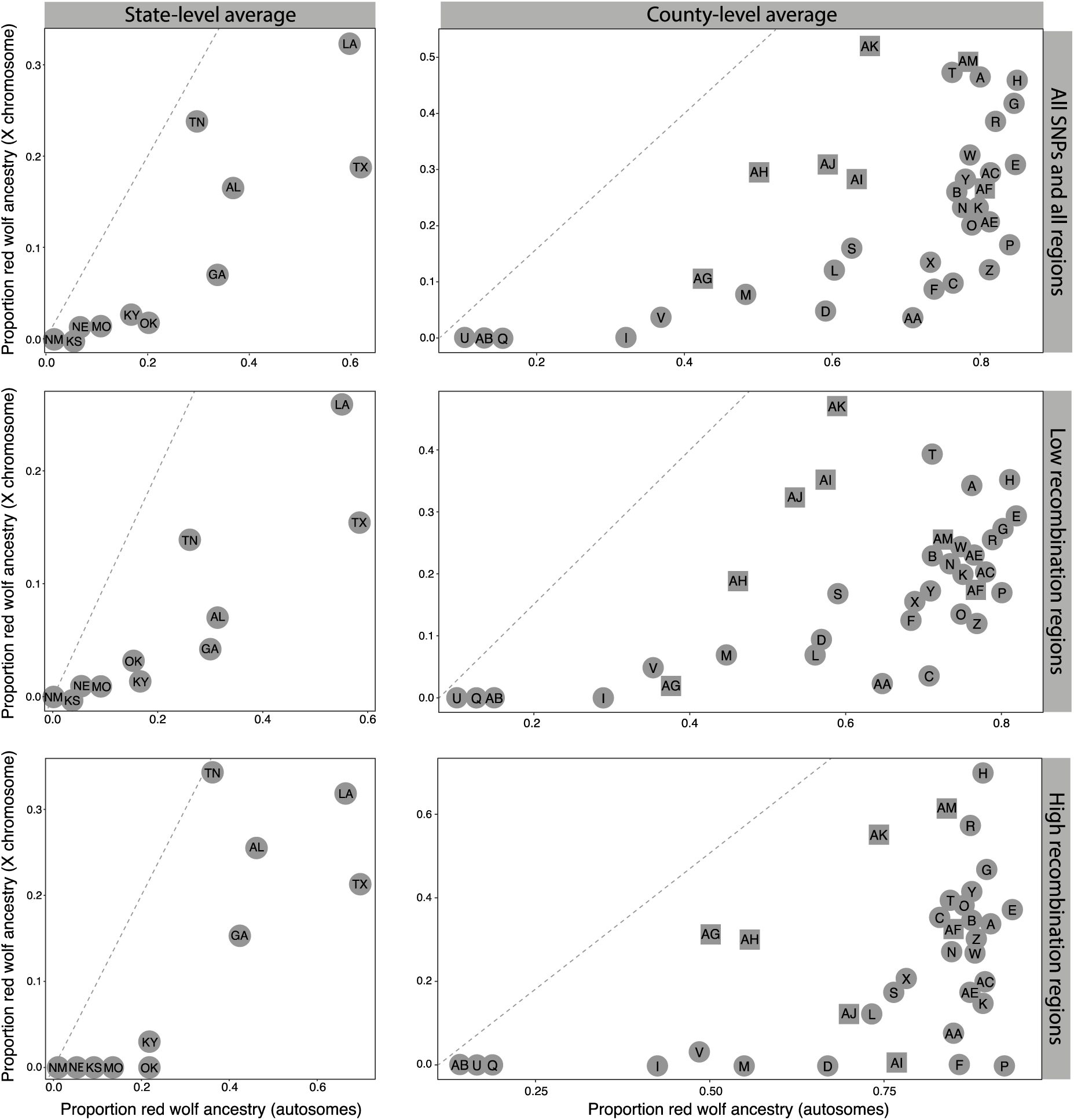
The average proportion of red wolf ancestry at the state- and county level (squares: Louisiana counties; circles: Texas counties) for 138 canids assigned using three reference populations for the 120K SNP set partitioned for all SNPs (top), low recombination regions (middle), and high recombination regions (bottom). See Table S3 for individual ancestry assignments to each of the reference populations. County abbreviations are as described in Figure 1B. (State abbreviations: AL, Alabama; GA, Georgia; KS, Kansas; KY, Kentucky; LA, Louisiana; MO, Missouri; NE, Nebraska; NM, New Mexico; OK, Oklahoma; TN, Tennessee; TX, Texas)

The distribution of red wolf ancestry switches and block sizes allowed us to estimate the timing at which red wolf alleles appeared in the ancestral genomes either through direct gene flow events or, more likely, through the dispersal of red wolf alleles through dispersing admixed coyotes. We used the average of two generation time estimates, the commonly estimated value of 4 years/generation and an estimate of 2 years/generation to account for scenarios in which a fraction of canids breed in their first year of life (Miller et al. 2003; Sacks 2005; vonHoldt et al. 2008; Hedrick et al. 2014; Albers et al. 2016; Mech et al. 2016; Kilgo et al. 2017). Autosomal red wolf admixture in coyotes was oldest in Louisiana (97 years ago, ya), whereas Tennessee coyotes carried the oldest X-linked red wolf ancestry (114ya) (Table S4A, Fig. 3). Coyotes within the historic range of the red wolf (Alabama, Georgia, Kentucky, Louisiana, Missouri, Oklahoma, Tennessee, Texas) acquired their red wolf ancestry on average 50ya (autosomal=53ya, X=54ya), which was significantly older than the red wolf ancestry found within coyotes of their own historic range (Kansas, Nebraska, New Mexico: autosomal=13ya, X=none, 1-tailed t-test *p*=0.0008). County-level time estimates further revealed the oldest autosomal red wolf admixture events within Louisiana where the last wild red wolves were documented (Plaquemines=234ya, Ouachita=121ya, Vermilion=98ya), relative to counties in western or southern Texas (e.g. Gaines=12ya, Irion=10ya, La Salle=26ya, Potter/Randall=15) (Table S4A). The X chromosome revealed much older red wolf ancestry in southeastern Texas (Austin=143ya, Bastrop=132ya, Chambers=117ya, Williamson=947ya) compared to Louisiana, which had much less X-linked red wolf ancestry detected (Table S4A), suggesting that the center of the historic red wolf population was likely in southeastern Texas.

**Figure 3.**
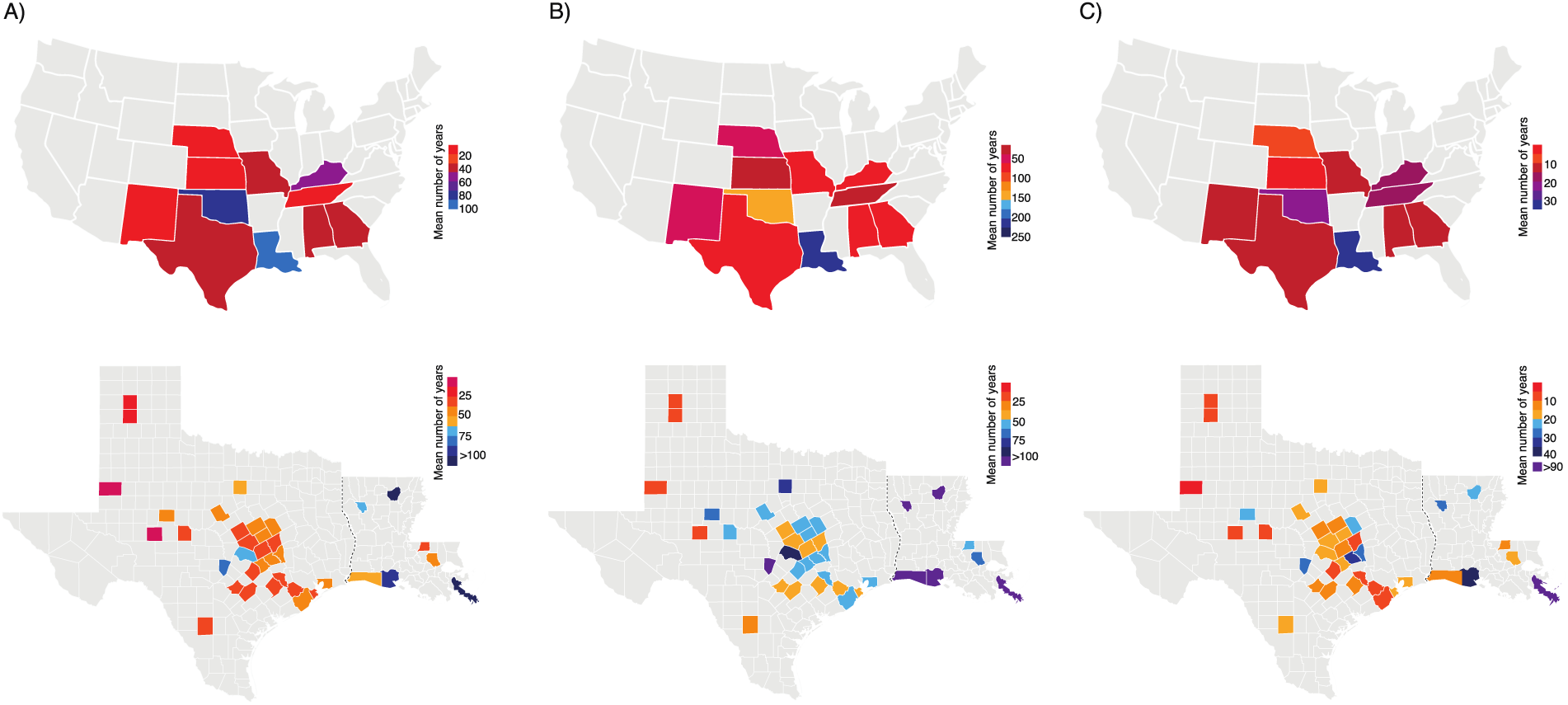
Average number of years since appearance of red wolf alleles across the autosomes for each state (top) and county (bottom) for 138 canids with inferred ancestry across **A)** the entire genome (120K SNPs), **B)** autosomal regions of low or **C)** high recombination rates.

### Recombination rates and ancestry

We partitioned the 120K SNP set into high and low recombination regions (<0.5cM/Mb or >2cM/Mb, respectively; Wong et al. 2010) to explore the impact of recombinational variation on admixture and demographic inference. We annotated 13,903 and 47,779 autosomal SNP loci as within regions of high and low recombination rates, respectively. Similarly, 430 and 1,297 X-linked SNPs were in regions of high and low recombination. Red wolf ancestry proportions were highest in coyotes from Texas and Louisiana across regions (autosomal, X average proportions: Texas=64.3%, 18.5%; Louisiana=60.9%, 28.8%) (Table S5; Fig. S10). There was a clear enrichment in red wolf ancestry in coyote autosomal regions with the highest recombination rates, and an enrichment of coyote ancestry in low recombining autosomal regions (Fig. S10). The X chromosome showed a similar trend with greater variance. Homozygous red wolf ancestry blocks were largest in autosomal regions of low recombination (average high=157Kb, low=779Kb) (Table S6). When we surveyed the X chromosome, regions of high recombination rates carried the longest tracks of homozygous red wolf ancestry (average high=230Kb, low=78Kb), with males carrying longer blocks than females (average high female=64Kb, male=92Kb, *p*>0.05) (Table S5). Coyotes from Alabama carried the longest tracks of homozygous red wolf ancestry regardless of recombination rate (low, autosome=1.2Mb, X=617Kb; high, autosome=718Kb, X=2.7Mb), followed by Tennessee, Louisiana, Texas and Georgia.

We found significantly older red wolf ancestry within low recombining chromosomal regions of coyotes within the historic range of the coyote (autosomal=92ya, X=673ya) compared to recently expanded southeastern populations (autosomal=24ya, X=no data; 1-tailed t-test *p*=0.0037) (Tables S4B,C; Fig. 2B,C). We found similar significant trends within chromosomal regions of high recombination rates (Historic: autosomal=18ya, X=32ya; Expansion: autosomal=6ya, X=no data, *p*=0.0146), which followed the expectations that low recombination regions carry signals of older demography where regions of high recombination likely display increased introgression. Louisiana coyote autosomes carried the oldest red wolf ancestry (Low=202ya, High=35ya), whereas Kansas carried the most recent (Low=11ya, High=2ya). Oldest red wolf autosomal ancestries were located within Louisiana parishes (Low=67-549ya, High=13-98ya), with Texas coyotes carrying the oldest X-linked red wolf ancestries (Low=42-1203ya, High=18-108ya). These events are most parsimoniously explained by the movement of red wolf ancestry through dispersal of admixed coyotes, especially given the declaration of the red wolf as extinct in the wild by 1980 (reviewed by Phillips et al. 2003).

## Discussion

In this study, we clearly demonstrate the persistence of red wolf genetic ancestry in admixed coyotes of the southern United States and that red wolf ancestry has remained relatively spatially restricted to the last regions that red wolves thrived. Although coyotes are a genetically diverse lineage with an apparent lack of local structure (Heppenheimer et al. 2018b,c), we found a strong increasing east-to-west geographic gradient of red wolf ancestry present in coyote genomes. This ancestry appears to derive predominantly from introgression rather than ILS, with the highest amounts of autosomal red wolf ancestry (∼60%) found in coyotes from southeastern Texas and southwestern Louisiana. We estimated that this ancestry was acquired up to 97 years ago through genetic exchange between coyotes and historic red wolf populations. However, we noted a depletion of red wolf ancestry on the coyote X chromosome relative to the autosomes, which carried the lowest proportions observed (∼14%), with the oldest red wolf content in a Tennessee coyote estimated to have been acquired 114 years ago.

These results advance our understanding of canid lineage evolution and hybridization, and have important implications for red wolf conservation. First, we present evidence that genetic exchange in the past century facilitated the introgression of red wolf alleles into the coyote populations of the Gulf coast, and red wolf ancestry had remained restricted to this region until fairly recently. This timeframe is consistent with the loss of the wild red wolf in the last century, where the detected geographic hot spot for cryptic red wolf ancestry was found in Louisiana and Texas, the last footholds of wild red wolves (Carley 1962; Paradiso 1968; Paradiso and Nowak 1971; Riley and McBride 1972; Carley 1975; MacCarley and Carley 1979). Concurrently with the decline of red wolves in this region due to habitat loss and persecution, coyotes expanded out of their historic range and encountered red wolves along their southeastern expansion front (Riley & McBride 1972; Nowak 1979; Parker 1995, Heppenheimer et al. 2018b; Hody and Kays 2018). Coyotes are a highly mobile species, often dispersing >100km (Harrison 1992; Sacks et al. 2005). However, our results indicate that only in the past two decades have introgressed red wolf alleles expanded north and westward in Texas. This observation does not appear to be attributed to a tension zone model, where a hybrid zone is maintained through the balance of ongoing hybridization and counter-selection within the zone (Barton and Hewitt 1985). Specifically, our data supports that the zone appears to be expanding rather than being constrained.

Here, we present genetic evidence that the red wolf is indeed a distinct species, supported through the identification of putative barrier or resistance loci, primarily on the X chromosome, but also on centromeric regions of some autosomes, a characteristic typically found in distinct species (Haasl and Payseur 2016; Harrison and Larson 2016). Although the red wolf has been considered a distinct species due to their unique ecology, morphology, and behavior (Waples et al. 2018), genetic and genomic distinctions have proved less well defined (e.g. vonHoldt et al. 2011, 2016a). We found low miscibility of the red wolf and coyote genomes, as evidenced by the paucity of red wolf ancestry on the coyote X chromosome. Further, we described a steep geographic cline of red wolf ancestry in the geographic region where the last wild red wolves were captured in the 1970s. The expansion of red wolf ancestry into coyote genomes has remained spatially constrained around these habitats. We thus suggest that these genomes have failed to “congeal” due to the X chromosome’s reduced capacity for introgression, where the entire X chromosome may in actuality act as a single super-locus (Turner 1967; Bierne et al. 2011). We propose that this lack of permeability has played a role in the divergence between coyotes and red wolves (Waples et al. 2018).

The hemizygous nature of sex chromosomes results in their reduced effective population size (Ne), accelerating the rate of evolution which can lead to the accumulation of loci involved with reproductive isolation (Kitano et al. 2009; Goldberg and Rosenberg 2015; Van Belleghem et al. 2018). Divergent evolutionary histories are common when inferred from sex chromosomes relative to the autosomes. This has been attributed to enhanced introgression barriers, which are suspected to result from the greater accumulation of incompatibilities that ultimately impact reproduction (the “large X-effect”) (Coyne and Orr 1989; Masly and Presgraves 2007; Garrigan et al. 2012; Martin et al. 2013; Sankararaman et al. 2014). Empirical evidence support supports these predictions, showing the X chromosome generally carries a reduced level of introgression (Payseur and Nachman 2005; Larson et al. 2014; Fontaine et al. 2015; Li et al. 2019; Fraïsse and Sachdeva 2021). This differential degree of X-linked ancestry is derived from the theory that in a population with equal males and females, two-thirds of the X chromosome are found in the females while one-third are in males (or the Z chromosome in the ZW systems). However, this linear assumption is violated when there are inequal contributions across the sexes in an ongoing-admixture model (Goldberg and Rosenberg 2015).

Similar introgression patterns have been well described in the genome of modern-day humans. Although humans and Neanderthals recently diverged (∼400,000-470,000 years ago, Harris and Nielsen 2016), the modern-day human X chromosome is depleted of Neanderthal ancestry (Sankararaman et al. 2014; Dutheil et al. 2015). The inference has been interpreted with respect to putative sterility and other reproductive barriers that prevent introgression, although this has been disputed given other phenomena could explain these patterns, such as sex-biased hybridization, stronger selection on the hemizygous nature of the sex chromosome, and differences in recombination rates (Hammer et al. 2010; Veeramah et al. 2014; Harris and Nielsen 2016; Juric et al. 2016). Long-term persistent of exogenous ancestry is expected to be shaped by the strength of selection and fitness consequences (Harris and Nielsen 2016). Reports of reduced X-linked ancestry suggest a role for recessive deleterious mutations as an isolating mechanism, in additional to strong selective sweeps (Dutheil et al. 2015; Harris and Nielsen 2016). Although previous studies indicated coyotes and red wolves have a more recent divergence (55,000-117,000 years ago, vonHoldt et al. 2016a), these estimates are likely confounded by recent and ongoing introgression because they did not account for local recombination rates. Indeed, a recent study showed that low recombinant regions of the X chromosome produced divergence time estimates between coyote and red wolf that are 20 times older than those derived from autosomal comparisons (Chafin et al. 2020). Such findings are consistent with those presented here that report an enrichment of reproductive isolation loci that drive the strong depletion of red wolf ancestry across the X chromosome.

Our study builds upon past efforts by factoring local recombination rates into our exploration of genealogies across divergent segments of chromosomes. As expected, we found that red wolf ancestry is highest (and younger) in regions of high recombination rates, consistent with theoretical expectations (Geraldes et al. 2011; Schumer et al. 2018; Seixas et al. 2018; Stevison and McGaugh 2020). In contrast, coyote X chromosomes show a paucity of red wolf ancestry, which we hypothesize is due to effects of interference across large regions with extremely reduced recombination rates, and parallels results in other mammalian systems (Carneiro et al. 2010; Li et al. 2019). These findings are expected following reduced the genomic permeability of barrier loci, consistent with our results that regions devoid of red wolf ancestry were estimated to be much older than those containing introgressed variation.

This study also has provided evidence that the coyote is a model system in which we can explore the dynamics and consequences of “hybridization load” in admixed individuals (Bierne et al. 2002; Juric et al. 2016; Harris and Nielsen 2016). The process of multi-generational admixture can produce a diversity of phenotypes due to recombinant genotypes, with an expectation that some of those phenotypes have higher relative fitness as they are shaped by natural selection (Barton 1979). For example, the human genome contains several archaic hominid introgressed regions with known fitness impacts, often an aspect of the yet nascent field of personalized medicine (Dolgova and Lao 2018; Durvasula and Sankararaman 2020; Rotival and Quintana-Murci 2020). The impact of specific introgressed regions and variants in coyote remain unknown, where theory suggests that fitness consequences of admixed genomes occur predominantly through the increased genetic load of weakly deleterious alleles that entered the genome from the parental species with a smaller effective size (i.e. red wolf). Several studies have uncovered genomic evidence of adaptive introgression in wild and domesticated species (e.g. Suarez-Gonzalez et al. 2016; Barbato et al. 2017; Figueiró et al. 2017; Jones et al. 2018; Schweizer et al. 2018; Burgarella et al. 2019; Oziolor et al. 2019; Sotola et al. 2019). Such an effort would likely inform both evolutionary theory as well as conservation efforts of red wolves, with respect to fitness considerations and species persistence.

Lastly, there are several implications for the future of red wolves. First, we have provided evidence that coyotes continue to be a significant reservoir of ghost and SSP lineage red wolf ancestry, a lineage that continues to become more inbred through time. Such introgression appears to have been a frequent event for southern coyotes during the past century. Future full genome sequencing efforts will allow us to more deeply explore the timing and genomic impact of introgression, including the identification of selection for and against red wolf alleles in specific genomic regions. Further, identifying and dating ancestry hot-spots in the historic red wolf range can help locate geographic regions that are ‘good’ at retaining red wolf characteristics and assist site selection for future red wolf reintroduction and recovery efforts. The longer-term viability of the red wolf relies heavily upon the potential for the SSP to incorporate novel and variable red wolf genomic content. We promote a species conservation design similar to that of de-introgression (Amador et al. 2014). Here, our suggestion is to enrich the SSP population for red wolf ghost genetics by careful breeding plans that incorporate controlled interbreeding of red wolves with coyotes that have high red wolf genomic content. Such a mechanism promotes the immediate transfer of the ghost genotypes once thought to be extinct in an effort to recover the native genetic background of the red wolf. Controlled breeding strategies for de-introgression or de-extinction that utilize hybrids have been proposed in other systems (Quinzin et al. 2019), as has the utilization of genome editing tools (Johnson et al. 2016), where these bold conservation efforts may become necessary in an era of global change. A de-introgression plan requires extensive screening of individuals for ancestry inference, sampling historic red wolf specimens to better characterize red wolf genetics prior to the 1970s, careful controlled breeding design, and subsequent promotion to the SSP breeding program. Although de-introgression is likely to be more successful when exogenous ancestry is limited and recent (Amador et al. 2013), the approach used here would provide a means for implementing an innovative recovery plan for the imperiled red wolf.

## Methods

### Sample Collection and DNA Extraction

We obtained 310 blood or tissue samples from state management programs, government organizations archives, or from museum archives. All samples collected have a known U.S. state of origin; many of which also have known county (Fig. 1A,B, Table S1). Samples were collected between 1987 and 2020. Reference populations are based on previous genome-wide studies that have identified populations of little to no admixture, as well as incorporating pre-expansion demographics (Heppenheimer et al. 2018a,d; Heppenheimer et al. 2020). Domestic dogs (*C. familiaris*) from North America were additionally included to address the possibility of recent hybridization, although little evidence suggests they have contributed towards wild canid ancestry (vonHoldt et al. 2011; Heppenheimer et al. 2018d). Reference red wolves were selected as captive individuals that genetically represented the red wolf founders with low pedigree inbreeding values (0.075-0.097). Further, we are cognizant that the demographic histories of the eastern wolf and gray wolves of the Great Lakes region is complex due to admixture and gene flow, which results in these populations known to carry a detectable, and sometimes substantial, amount of non-gray wolf ancestry (Heppenheimer et al. 2018d). We obtained high molecular weight genomic DNA from either the Qiagen DNeasy Blood and Tissue Kit or the BioSprint 96 DNA Blood Kit performed on the Thermo Scientific KingFisher Flex Purification platform and following manufacturer’s blood or tissue protocol for mammals. We quantified DNA concentration using either the PicoGreen or Qubit 2.0 fluorometry systems, and subsequently standardized DNA samples to 5ng/µL.

### RAD sequencing and bioinformatic processing

We prepared 205 genomic libraries for restriction-site associated DNA sequencing (RADseq) following a modified protocol (Ali et al. 2015). We digested genomic DNA with *SbfI* and ligated a unique 8-bp barcoded biotinylated adapter to the resulting fragments. We then pooled equal amounts of 48 DNA samples followed by random shearing to 400bp in a Covaris LE220. We then enriched the sheared DNA pools for adapter ligated fragments using a Dynabeads M-280 streptavidin binding assay. We prepared these enriched pools using the NEBnext Ultra II DNA Library Prep Kit for Illumina NovaSeq 2−150nt sequencing at Princeton University’s Lewis Sigler Genomics Institute core facility. We selected fragments 300-400bp in size with Agencourt AMPure XP magnetic beads, which were also used for library purification. Sequence data were first processed to identify which read (forward or reverse) contained the unique barcode and the remnant *sbfI* cut site. Using a custom perl script, these reads were then retained in a single file, the matching read pairs that lacked the cut site were curated in a separate file, and all remaining reads were discarded. We then processed these reads in *STACKS* v2. We first demultiplexed using *process_radtags* using a 2 bp mismatch for barcode rescue and retained reads with a quality score ≥10. We next removed PCR duplicates with the paired-end sequencing filtering option in *clone*_*filter* and then mapped to the dog genome CanFam3.1 assembly (Lindblad-Toh et al. 2005) using *STAMPY* v1.0.21 (Lunter and Goodson 2011). We additionally filtered mapped reads for a minimum MAPQ of 96 and converted to bam format in *Samtools* v0.1.18 (Li et al. 2009). We included 105 publicly available canid samples already mapped to the same reference genome assembly following these methods (Table S1).

We completed SNP discovery using all samples with a minimum of 100,000 mapped reads to obtain a catalogue of all polymorphic sites possible. We implemented the *gstacks* and *populations* modules in *STACKS* v2 following the recommended pipeline for data mapped to a reference genome (Catchen et al. 2013; Rochette et al. 2019). We increased the minimum significance threshold in *gstacks* to require more stringent confidence needed to identify a polymorphic site using the marukilow model (flags --*vt-alpha* and --*gt-alpha*, *p*=0.01). We then selected a subset of individual samples with a minimum of 200,000 mapped reads for comparable geographic and lineage representation in the *populations* module. We reported all SNPs discovered per locus (opted against using the *populations* flag --*write_single_snp*) as ancestry inference is best with high density data. We established a *gstacks* catalogue of 2,799,665 variants in *STACKS* v2 (Catchen et al. 2013; Rochette et al. 2019), of which 94,277 SNPs were retained after filtering for a minimum of 3% minor allele frequency (MAF) and 20% missingness. We further filtered to retain sites with a MAF>3% and allowed up to 70% genotyping rate per locus (flag --*geno* 0.3) for the initial filtering step in *PLINK* v1.90b3i (Chang et al. 2015). To determine if any individual sample was missing a significant proportion of genotypes, we assessed missingness using the *PLINK* function --missing. We excluded 17 samples that had >40% missing genotype data (Table S1) and repeated the *stacks populations* module for recalling SNP genotype across 293 canids at 2,759,705 SNP loci. This sample set included all reference species and populations in addition to the wide geographic representation across the southern United States. We retained 120,621 after filtering for 3% MAF and 20% missingness (referred to as 120K SNP set) with an average density of one SNP every 19Kb. For demographic analyses of neutral aspects of genetic structure and diversity estimates, we constructed a “*statistically neutral and unlinked*” dataset of SNPs by excluding sites within 50-SNP windows that exceeded genotype correlations of 0.5 (with the *PLINK* argument *--indep-pairwise* 50 5 0.5) and deviated from Hardy-Weinberg Equilibrium (HWE) with the argument --*hwe* 0.001. This resulted in a statistically neutral and unlinked dataset of 83,851 SNP loci (referred to as 83K SNP set) for demographic analyses (n SNPs autosomes=83,021 and X chromosome=830). The X chromosome was analyzed independently from the autosomes to ensure the inclusion of the signal driven by the single sex chromosome captured in our sequencing and mapping efforts.

### Sex inference

To bioinformatically infer the sex of canids, we re-mapped FASTQ files to the complete CanFam3.1 reference assembly, with the addition of the Y chromosome (GenBank KP091776.1; Li et al. 2013). We estimated the number of reads that covered each nucleotide on the Y chromosome per genome. We expected a distinct difference in the number of Y-linked reads that map for males versus females, with some variation expected for females due to reads that map to the Y chromosome within to the canine pseudoautosomal region (PAR) on the X chromosome (1bp-6.7Mb; Raudsepp et al. 2012). Several samples had field-based sex reported which were surveyed for accuracy (Table S1). From animals with known sex records, we estimated a 5.2% discordance rate in 96 canids analyzed, where three females and two males had a genomic sex inference that did not match field-based records (Table S1; Fig. S6). For the remaining 42 canids, we inferred 19 females and 23 males.

### Population structure analysis

To survey genetic clustering, we conducted a PCA with *flashPCA* (Abraham and Inouye 2014). We then employed a maximum likelihood clustering method to estimate the most likely number of genomic clusters (*K*) in *ADMIXTURE* v1.3. (Alexander et al. 2009) with the cross-validation flag to assess inter-*K* likelihoods.

### Assessment of incomplete lineage sorting and gene flow

Following from the aforementioned methods in *STACKS* v2 for cataloguing and discovering SNP variants, we also independently repeated these methods with a larger dataset that included 45 red fox (*Vulpes vulpes*) previously published, with at least 100,000 mapped sequence reads (SRA PRJNA510648) (DeCandia et al. 2019). We filtered in *VCFtools* v0.1.17 (Danecek et al. 2011) to remove loci with >10% missing data across all individuals, singletons and private doubletons, and individuals with >20% missing data. We identified loci that are fixed in all 45 of the red fox genomes, which are informative for the ancestral allele state. Any locus that was lacking data in any of the groups was also excluded from the analysis. For the *ABBA-BABA* inference of ILS or introgression, we implemented an allele-frequency approach for estimating *D* statistics in R using derived allele frequencies across the red fox as an outgroup, the SSP red wolf as P3, reference coyote as P2, and several permutations of coyotes in P1 (following Martin et al. 2013). The autosomes and the X chromosome were analyzed as a single dataset. We assessed statistical significance through a block jackknife approach with block sizes from 2-10Mb.

### Inference of canid ancestry

We inferred local ancestry of 138 coyotes with possible red wolf ancestry (n Alabama=5, Georgia=8, Kansas=2, Kentucky=5, Louisiana=10, Missouri=11, Nebraska=5, New Mexico=1, Oklahoma=5, Tennessee=3, Texas=83) with respect to three reference populations (coyote, red wolf, and gray wolf) (Table S3). We implemented a two-layer hidden Markov model in the program *ELAI* (Guan 2014). This approach first evaluates LD within and between the reference canid groups and returns a per-SNP allele dosage score, which estimates the most likely ancestry proportion (i.e. homozygous scores are 0 and 2, heterozygous score is 1). SNPs were automatically excluded if they were missing from one of the populations. We analyzed unphased SNPs that were unfiltered for genotype correlations or HWE. We defined the number of upper-layer clusters (-*C*) to 3 (i.e., red wolves and each reference coyote population) and the lower-layer clusters (-*c*) to 15 (5x the value of *C*, as recommended). To account for both uncertainty in the precise timing of admixture and increased complexity in admixture scenarios where only a single-pulse of admixture is unlikely, we analyzed four timepoints since admixture (-*mg*): 5, 10, 15, and 20 generations. We implemented *ELAI* three times serially for each admixture generation value with 30 EM steps and averaged results over all 12 independent analyses. We considered sites with allele dosage between 0.5 and 1.5 to be heterozygous and sites with allele dosage >1.5 to be homozygous. Sites with multiple and incompatible ancestries (e.g. three heterozygous states) were excluded from the ancestry block analyses. We required that ancestry blocks contain a minimum of two contiguous SNPs with the same ancestry assignment. We also assumed that all ancestry (via allele dosage) outside of the PAR on the X chromosome of males was haploid in ancestry (allele dosage>1).

### Estimating the timing of admixture

We counted the number of ancestry block identity switches per individual genome, with a focus on the genome-wide ancestry estimates of red wolf and gray wolf. Given the reduced representation focus on *Sbf1* cut sites and size selection step, the resulting blocks are likely inflated in size with a paucity of small block sizes. Although this design is incredibly useful for rapid genotyping of thousands of SNP loci across hundreds of genomes, the estimates are likely skewed towards more recent timing of admixture events given the density of markers. Following from Johnson et al. (2011), we then used the below equation to estimate the number of generations since admixture for diploid genomes:

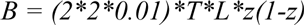

where *B* is the estimated number of ancestry switches, *T* is the number of generations since admixture, *L* is the total genome length (2085cM for autosomes and 111cM for the X chromosome; Wong et al. 2010), and *z* is the genome-wide ancestry proportion of either red wolf or gray wolf specific to autosomes or X chromosome.

### Recombination rate integration

To determine how recombination variation along the genome influences the patterns of admixture and structure, we integrated recombination rates from the autosomes and X chromosome (Wong et al. 2010), with all positions from CanFam2 lifted over to the CanFam3.1 reference genome assembly. Following from Li et al. (2019), we implemented a two-step approach for assigning recombination rates and smoothed averages. We first estimated recombination rates (cM/Mb) for each marker within 100Kb non-overlapping windows (step=100Kb) and used the female-based estimates for the X chromosome. To mitigate finer-scale variation in recombination rates and focus on broader scale rates, we smoothed recombination rate averages across 2Mb sliding windows. These resulting windows were then partitioned into either low or high recombination rate windows if their smoothed average was <0.5cM/Mb or >2cM/Mb, respectively. Each SNP locus contained within a high or low recombination window were parsed into separate datasets for analysis using the *intersect* function flag *-loj* of *BEDtools* v2.28.0. Similarly, as described above, we estimated admixture timing for high and low recombination blocks with their respective total centimorgan lengths analyzed (autosomes: low=820cM, high=697cM; X chromosome: low=8cM, high=37cM).

### Ethics and data availability

This work was conducted under the approved Princeton University IACUC protocol 1961A-13. RADseq BAM files sequenced in this study have been submitted to the NCBI BioProject database (https://www.ncbi.nlm.nih.gov/bioproject/) under accession number PRJNA684924. See Table S1 for references to additional public data included in this study.

## Acknowledgements

We graciously thank Sarah Hamer, Mike Pipas, Karin Saucedo, Steve Parker, Ron Wooten, Dianna Krejsa, Clint Harper, J Bennet, and Roland Kays for providing samples and contacts. We also thank Elizabeth Heppenheimer for advice regarding analytical methods. The findings and conclusions in this article are those of the authors and do not necessarily represent the views of the U.S. Fish and Wildlife Service.

## Author contributions

BV and WM designed the study; BV, KB, CS, LYR, SRF, and AS provided sample collections; AL assisted in genomic DNA purification; WM provided the dog Y chromosome sequence; BV conducted the analyses; WM and MA provided analytical feedback; all authors contributed towards the manuscript preparation.

---

**Supplemental Table S1.** Meta-data for each canid sample. (Abbreviations: F, female; M, male)

See file Supplemental_Table_S1.xlsx

**Supplemental Table S2.** Per partition (*K*) assignment probabilities for 293 canids genotyped at **A)** 83K and **B)** 830 X-linked neutral and statistically unlinked SNP loci; and 254 canids genotyped at **C)** 83K and **D)** 830 X-linked neutral and statistically unlinked SNP loci. (Abbreviations: SSP, Species Survival Plan)

See file Supplemental_Table_S2.xlsx

**Supplemental Table S3.** Ancestry switches, proportion, and blocks metrics for 138 individual canids across autosomes and the X chromosome. For timing, estimates are given for a 2-year and 4-year generation time. (Abbreviations: Auto, autosomes; X, X chromosome)

See file Supplemental_Table_S3.xlsx

**Supplemental Table S4.** Gene flow timing estimates for 138 individual canids across autosomes and the X chromosome **A)** genome-wide, **B)** in low recombination regions, and **C)** in high recombination regions. For timing, estimates are given for a 2-year and 4-year generation time. (Abbreviations: Auto, autosomes; X, X chromosome)

See file Supplemental_Table_S4.xlsx

**Supplemental Table S5.** Ancestry switches, proportion, and blocks metrics for 138 individual canids for low and high recombination blocks across the genome. For timing, estimates are given for a 2-year and 4-year generation time. (Abbreviations: Auto, autosomes; X, X chromosome)

See file Supplemental_Table_S5.xlsx

**Supplemental Figure S1.**
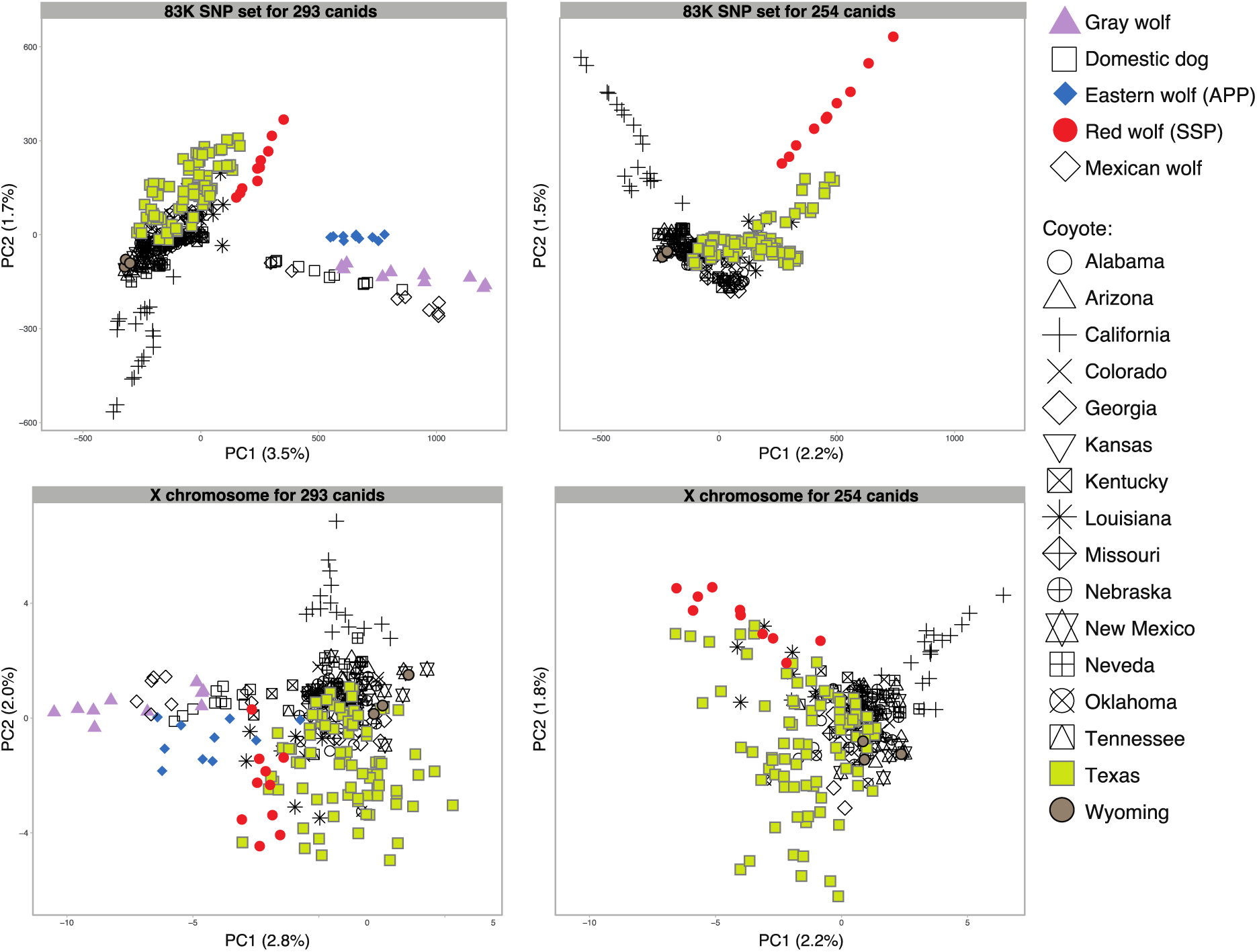
PCA of the 83K neutral and unlinked SNP set and separately for 830 SNPs on the X chromosome for 293 and 254 canids. See panel labels below, with each axes labeled with percent of variance explained.

**Supplemental Figure S2.**
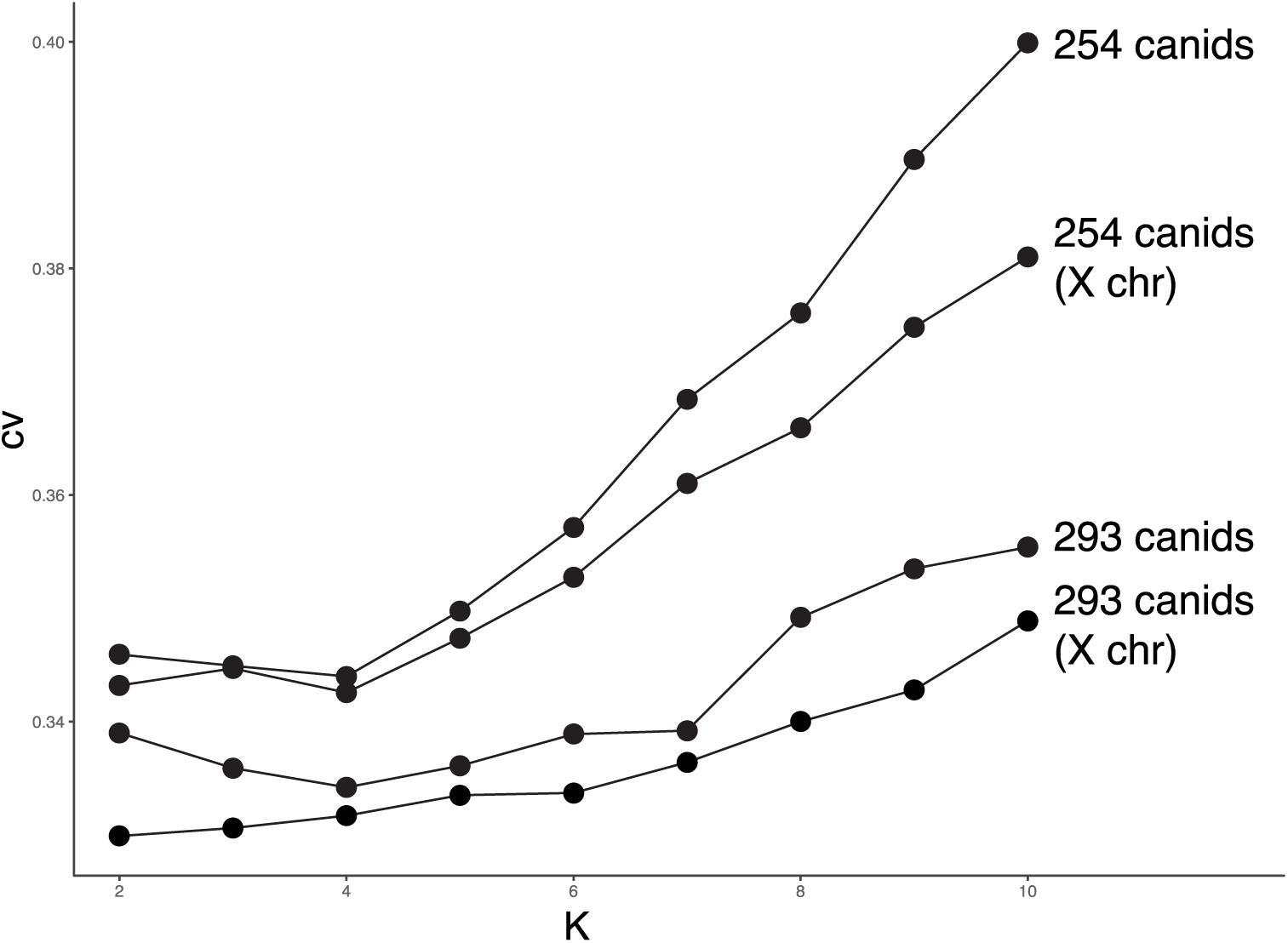
Cross-validation (cv) error rates from *Admixture* analysis of canids genotyped at unlinked neutral SNP loci. Lowest cv errors indicate the best fit partition, although multiple partitions are often informative and displayed. See main text for number of SNPs per sample set. (Abbreviations: chr, chromosome)

**Supplemental Figure S3.**
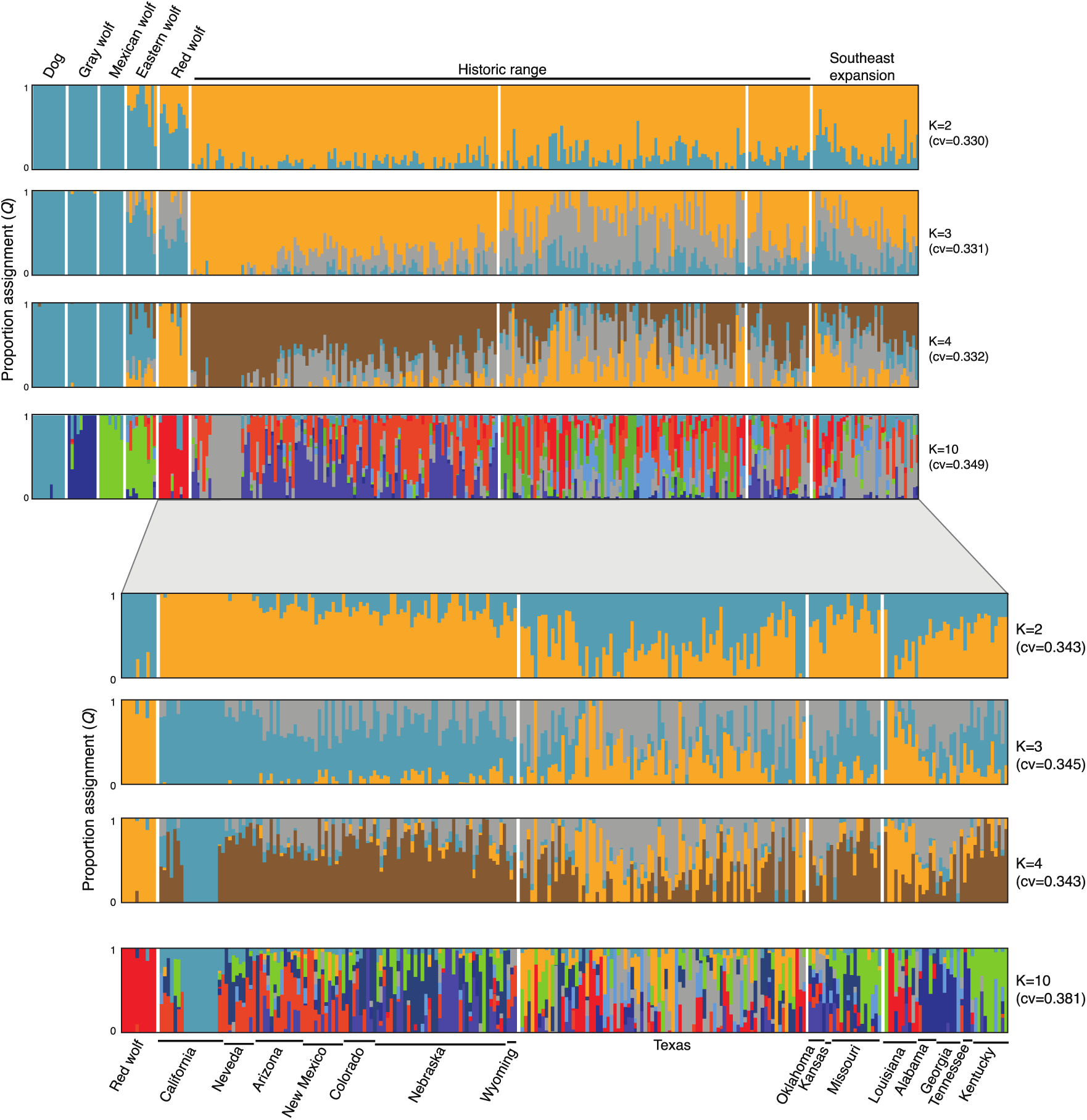
Admixture analyzes of 830 neutral and unlinked SNPs across the X chromosome for 293 and 254 canids (top and bottom panels, respectively). Solid bars above or below vertical bars indicate the state from which the samples originated.

**Supplemental Figure S4.**
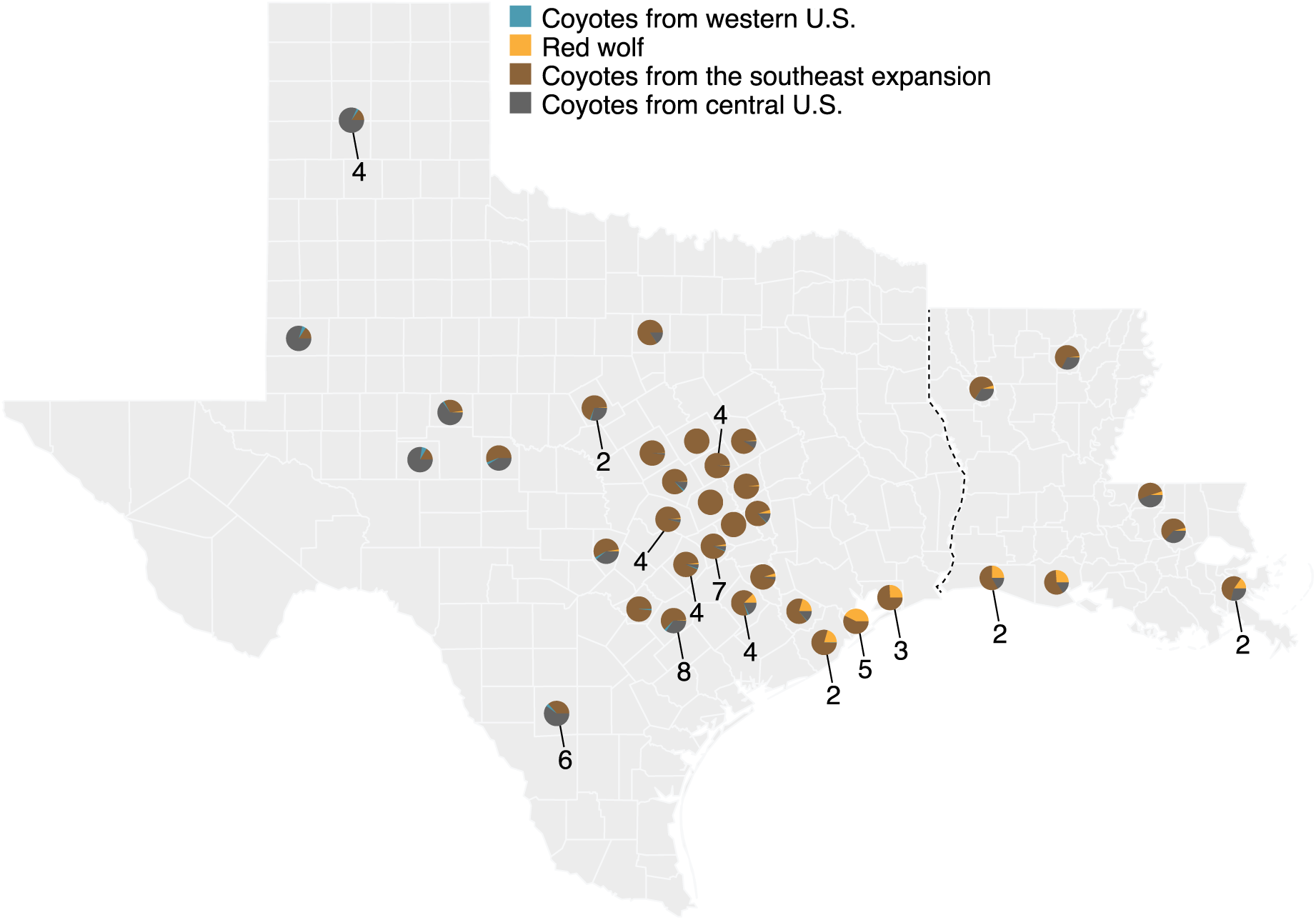
Averaged admixture proportions (*Q*) of 74 coyotes from Texas and nine coyotes from Louisiana with known county of origins at *K*=4 for 83K neutral and unlinked SNPs genotyped across the 254 sample set (excluded dogs, gray wolves, Mexican wolves, and eastern wolves). The legend indicates to which genetic cluster the population average proportions were assigned. Sample sizes per location are provided only when n>1. For detailed *Q* values, see Table S2C.

**Supplemental Figure S5.**
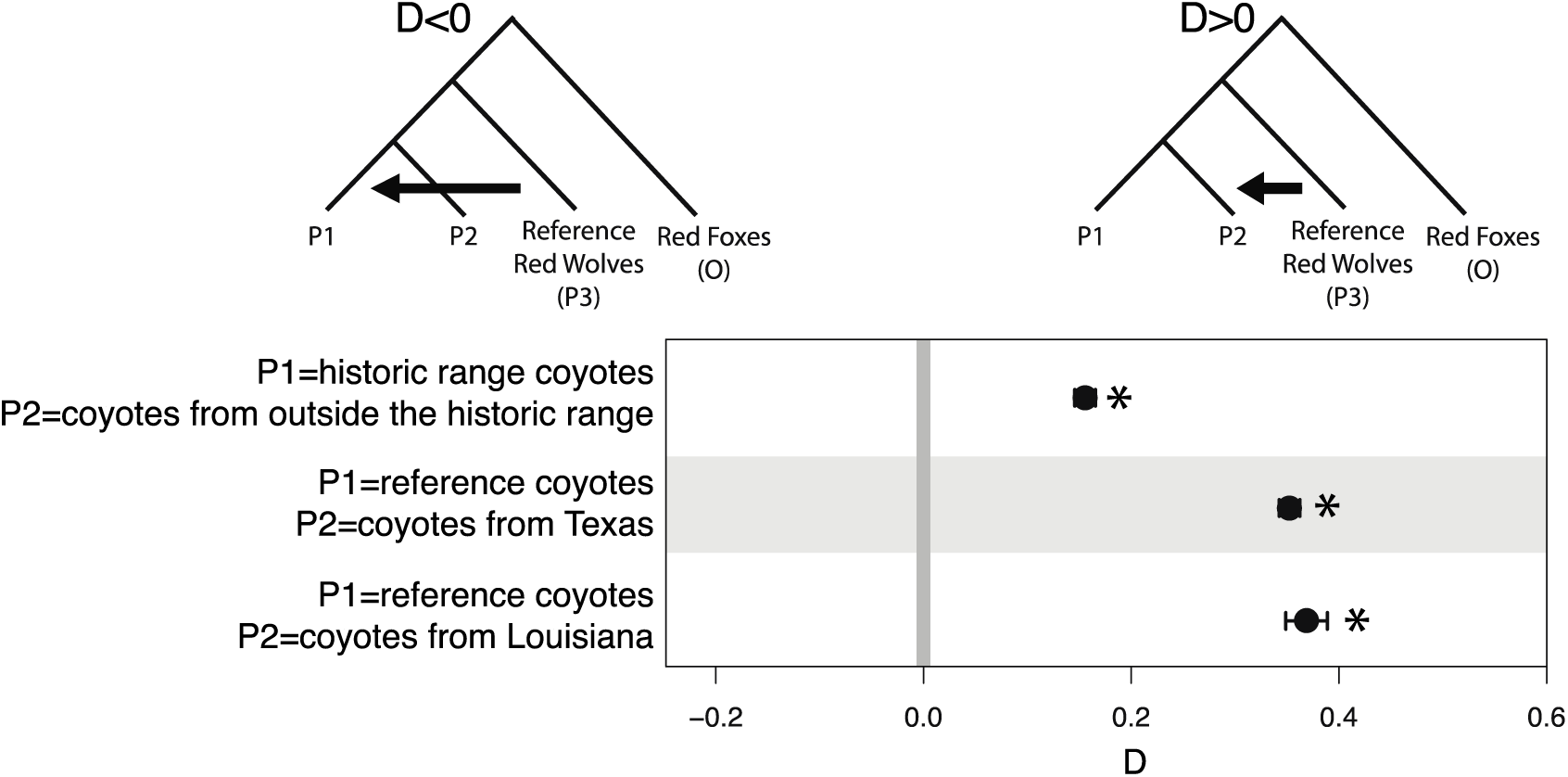
*D*-statistic testing for three P2 groups of coyotes. Standard error bars and significance (* *p*<0.05) were evaluated using jackknife resampling method with a block size of 5Mb. A value of *D<*0 would suggest incomplete lineage sorting, while *D*>0 values suggest introgression and gene flow. See Table 1 for details. (Abbreviations: O, outgroup)

**Supplemental Figure S6.**
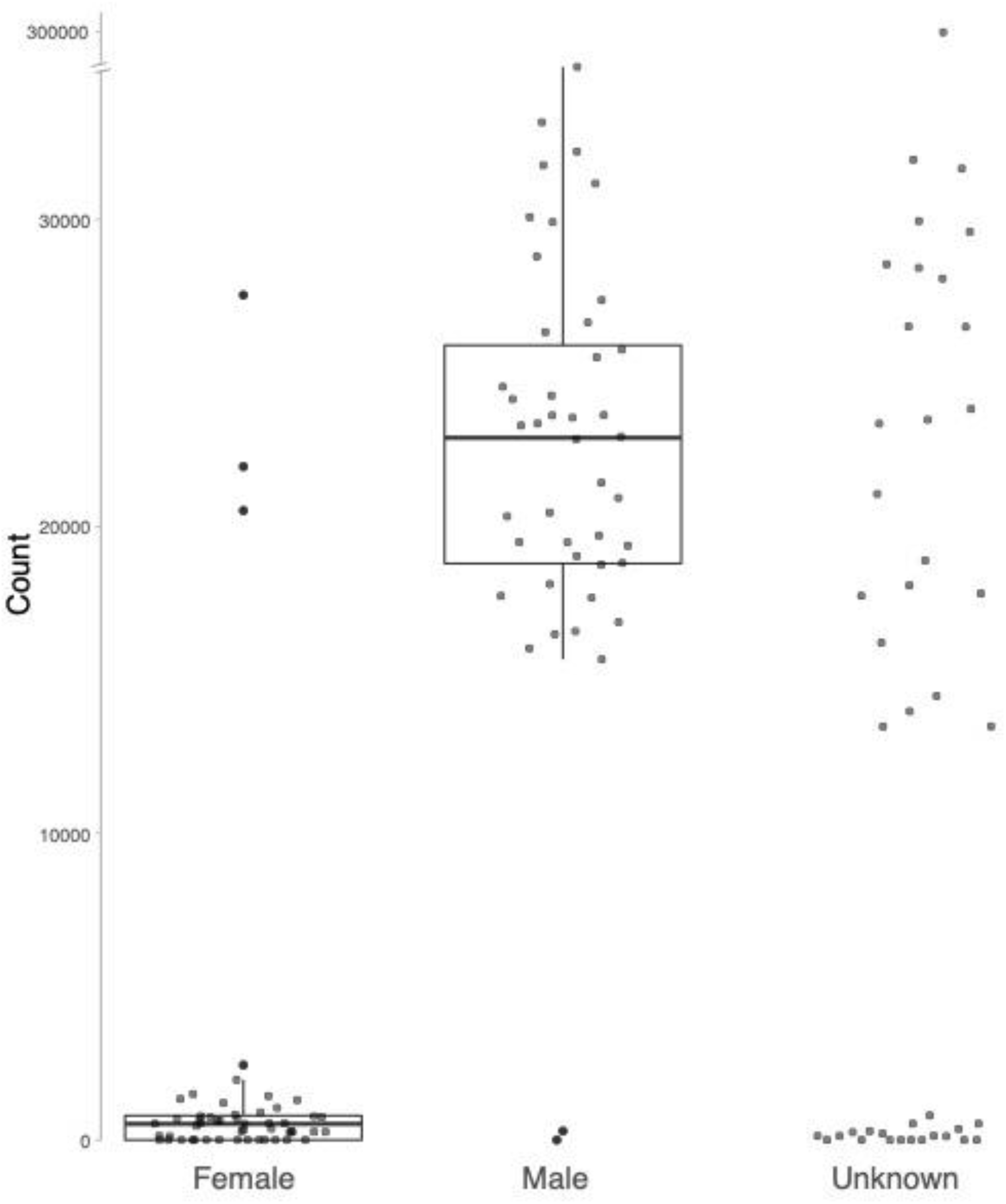
Distribution of the number of nucleotides (Y axis) mapped to the canine Y chromosome for bioinformatic sex inference across 138 canids targeted for ancestry inference. Of these canids, 96 had known sex documented from observation (42 canids were previously of unknown sex). See Table S1 for details.

**Supplemental Figure S7.**
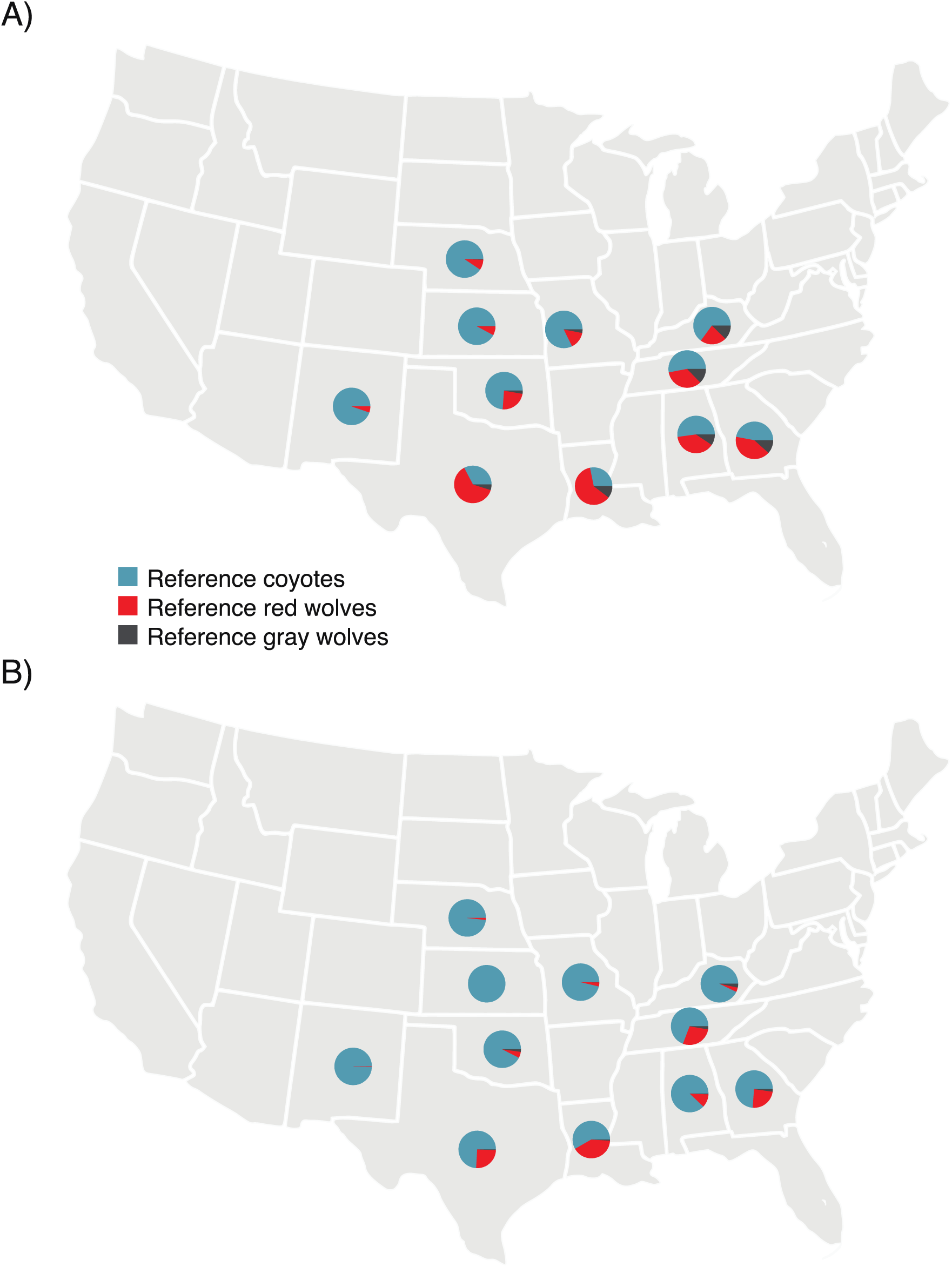
Per-state average ancestry inference for 138 canids from a three-ancestor model for **A)** autosomes and **B)** the X chromosome from the 120K SNP set. See Table S3 for individual ancestry proportions.

**Supplemental Figure S8.**
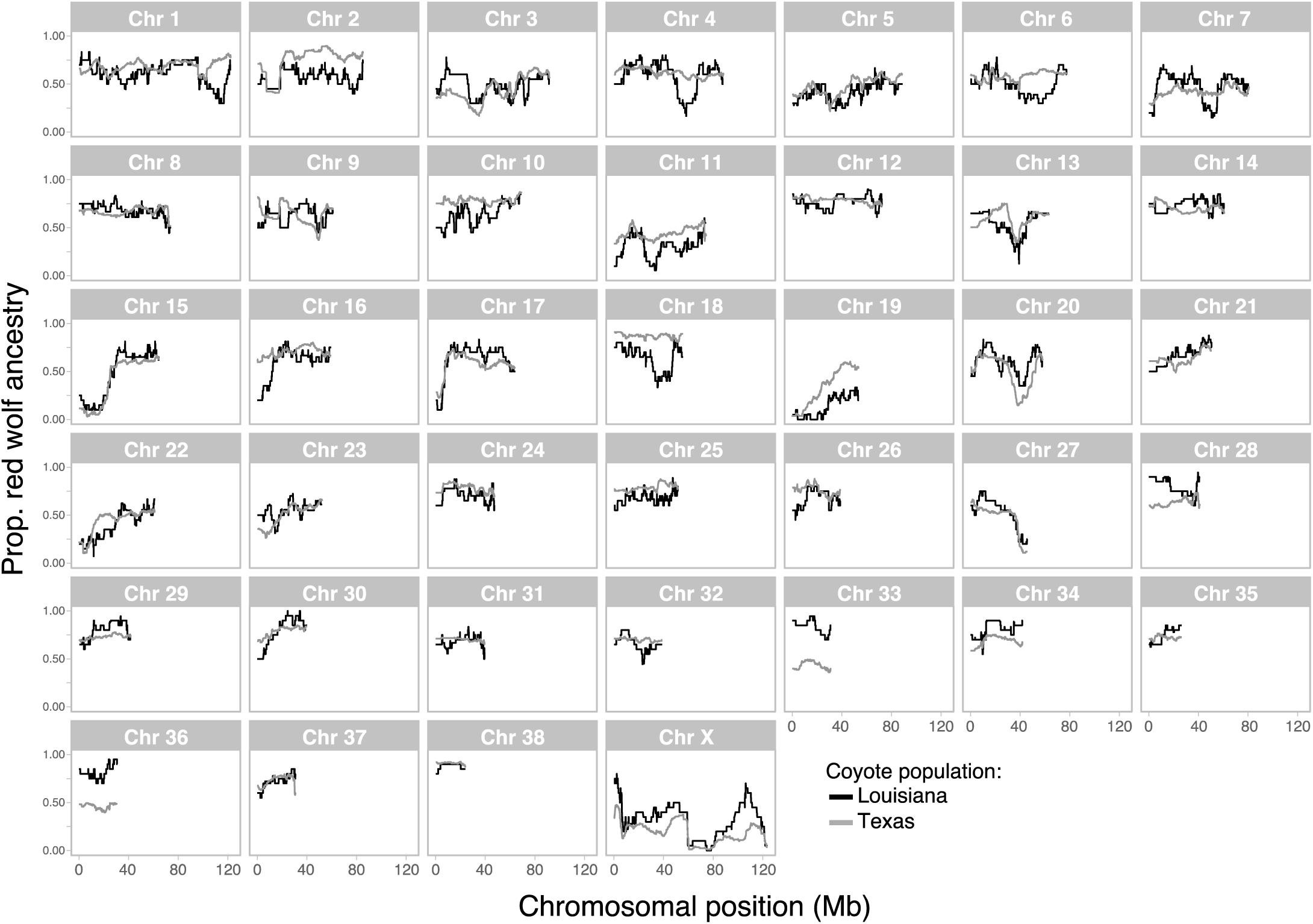
Per chromosome frequency of the proportion of red wolf ancestry across the putatively admixed coyote population of Louisiana (n=10) and Texas (n=83) genotyped at 120K SNP loci.

**Supplemental Figure S9.**
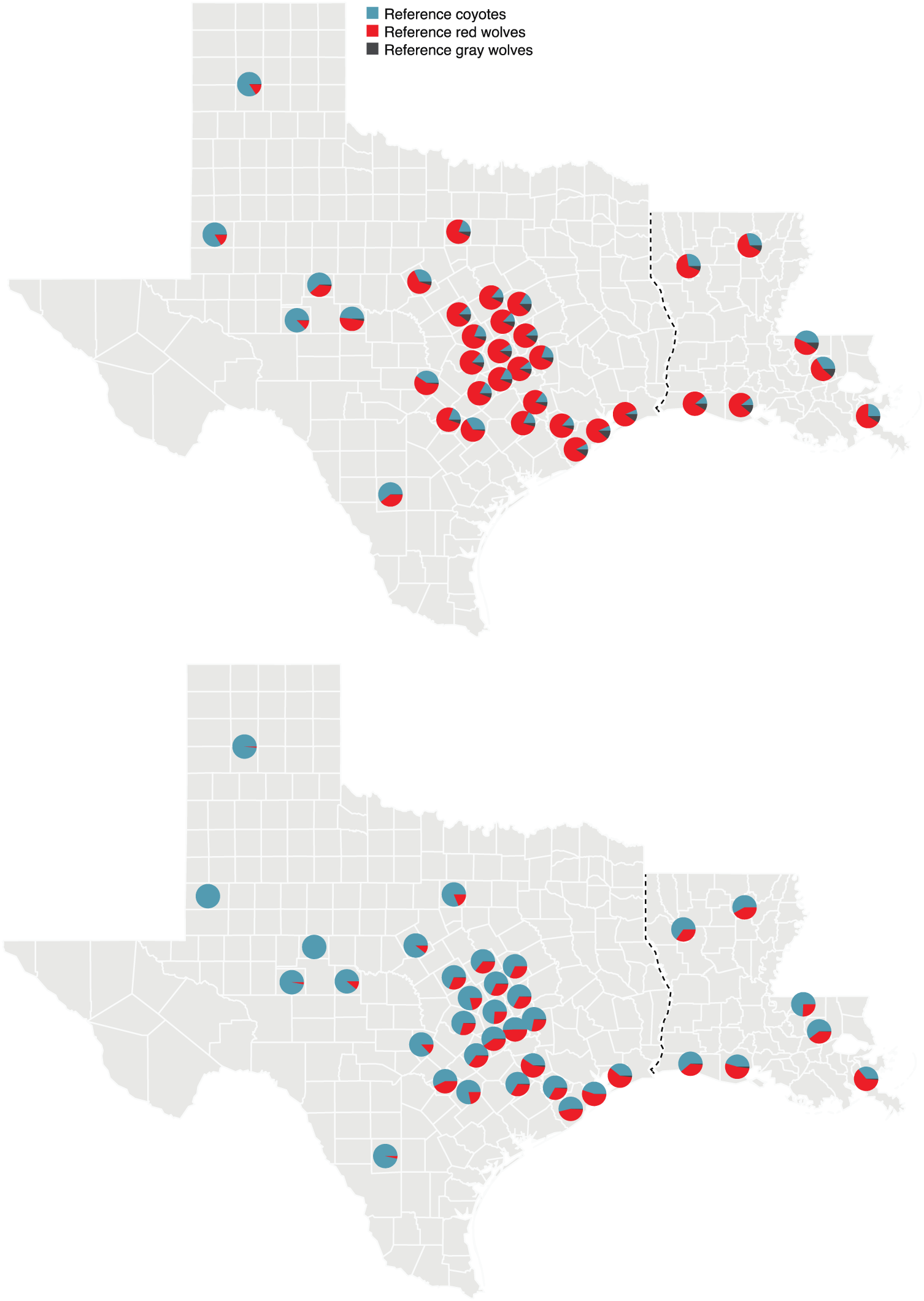
Per-county average autosomal (top) and X chromosome (bottom) ancestry inference for 138 canids from a three-ancestor model. See Table S3 for individual ancestry proportions.

**Supplemental Figure S10.**
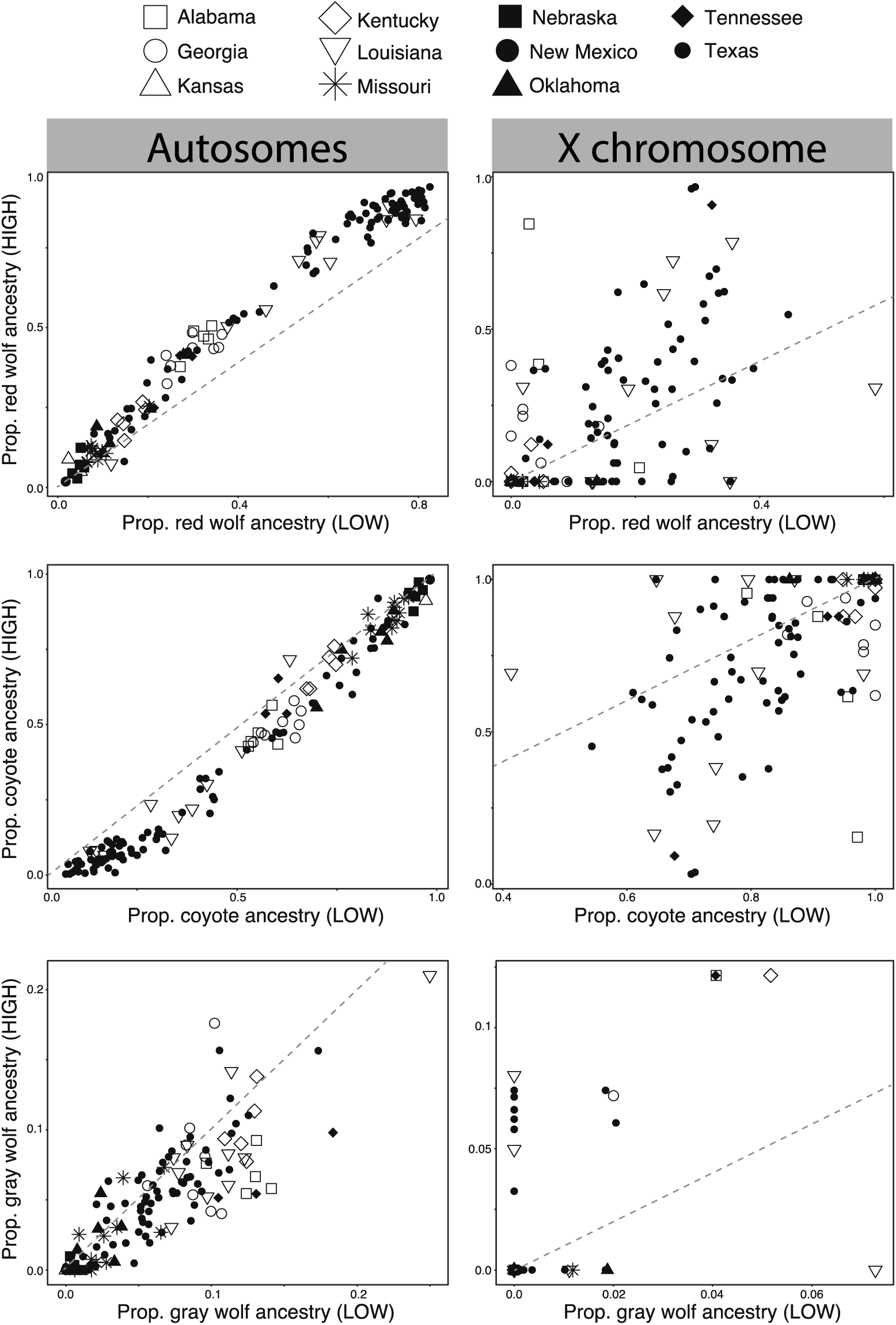
The average ancestry proportion for 138 canids assigned using three reference populations for autosomal (left) or X-linked (right) SNPs as a function of their recombination rates (low versus high). Dashed line indicates a 1:1 correspondence of ancestry across the two recombination rates. See Table S5 for details.

